# Gadolinium-free Magnetic Resonance Imaging of the Liver using an Oatp1-targeted Manganese(III) Porphyrin

**DOI:** 10.1101/2021.08.04.455144

**Authors:** Nivin N. Nyström, Hanlin Liu, Francisco M. Martinez, Xiao-an Zhang, Timothy J. Scholl, John A. Ronald

## Abstract

Controversy surrounding gadolinium-based contrast agents (GBCAs) have rendered their continued utility highly contentious, but the liver-specific GBCA Gd(III) ethoxybenzyl-diethylene triamine pentaacetic acid (Gd(III)-EOB-DTPA) remains in use because it provides unique diagnostic information that could not be obtained by any other means. To address the need for an alternative liver-specific MRI contrast agent, we synthesized Mn(III) 20-(4-ethoxyphenyl) porphyrin-5,10,15-tricarboxylate (Mn(III)TriCP-PhOEt), which exhibited significantly higher *r*_1_ relaxivity than Gd(III)-EOB-DTPA, and targeted organic anion-transporting polypeptide 1 (Oatp1) channels as a biomarker of hepatocyte viability. Mn(III)TriCP-PhoEt increased the *r*_1_ relaxation rate of cells expressing rodent *Oatp1a1* and human *Oatp1b3*, relative to control cells not expressing these liver channels. In mice, Mn(III)TriCP-PhoEt resulted in significant and specific increases in liver signal intensity on *T*_1_-weighted images, and significant decreases in liver *T*_1_ time relative to precontrast measurements. Our findings suggest that Mn(III)TriCP-PhOEt operates as a specific and sensitive MR contrast agent for *in vivo* liver imaging.

## INTRODUCTION

Magnetic resonance imaging (MRI) is widely applied for clinical diagnosis of disease by generating detailed high-resolution, 3-dimensional images of deep tissues in living subjects. Native image contrast is derived from the magnetic relaxation properties and distribution of abundant water protons (^1^H) in soft tissues. In addition, contrast agents can be used to shorten the longitudinal (*T*_1_) or transverse (*T*_2_) nuclear spin relaxation time of bulk water protons, and in turn, increase the contrast of tissues, vessels, cells or molecular events that would otherwise go undetected with non-contrast ^1^H MRI. In the clinic, gadolinium-based contrast agents (GBCAs) are administered in approximately 40% of patient scans tens of millions of times annually.^[1]^ GBCAs have provided essential diagnostic information that often cannot be obtained by other noninvasive modalities, making them highly beneficial.

No safety concerns existed with the use of GBCAs until 2006, when a seminal study^[2]^ linked Gd(III) exposure to the onset of a debilitating and sometimes fatal disease called nephrogenic systemic fibrosis (NSF), the first cases of which surfaced in 1997.^[3]^ With identification of severe renal impairment as a major risk factor for NSF, limiting access of GBCAs to patients with low glomerular filtration rates has reduced cases to single digits.^[4]^ More recently however, GBCAs have received scrutiny due to the discovery of long-term deposition in the central nervous system, liver, skin and bone of patients with otherwise healthy kidneys. This has been particularly evident in patients that received multiple doses of GBCAs where concentrations of retained Gd(III) in tissues rose above millimolar detection thresholds on MR images.^[5]^ Beyond the clinic, GBCAs are also emerging as environmental contaminants in aquatic ecosystems, especially in locations surrounding major medical hubs.^[6]^

Various approaches have been pursued to address concerns associated with GBCAs, including chelation therapies administered after image acquisition^[7]^ and artificial intelligence-based methods that create synthetic contrast-enhanced images,^[8]^ but while these are in early stages of development, Gd(III)-free contrast agents remain the most reliable method for generating contrast enhancement without GBCA-associated risks.^[1]^ Contrast enhancement on MRI can be achieved by various other compounds including manganese-based contrast agents (MBCAs), superparamagnetic iron oxide nanoparticles (SPIONs), exchangeable solute protons for chemical exchange saturation transfer (CEST) imaging, X-nuclei for non-proton MRI, and hyperpolarized imaging.^[9]^ Of these, MBCAs generate *T*_1_-weighted contrast with *in vivo* kinetics and biodistributions most like GBCAs, and do not require specialized pulse sequences, further postprocessing, or auxiliary hardware.^[9]^ Collectively, these aspects render MBCAs as attractive alternatives to GBCAs for contrast-enhanced MRI.

Notably, *in vitro* and animal studies have demonstrated that linear GBCAs in particular exhibit increased risks due to decreased chelation stabilities and increased tissue retention rates relative to macrocyclic GBCAs.^[10]^ In 2017, the U.S. Food and Drug Administration (FDA) announced a class warning for all GBCAs^[11]^ and the European Medicines Agency (EMA) followed^[12]^ by suspending marketing authorizations for three of the five GBCAs comprised of linear chelators as the “benefit-risk balance [was] no longer favorable.” During these announcements however, Gd(III) ethoxybenzyl-diethylene triamine pentaacetic acid (Gd(III)-EOB-DTPA; Gadoxetic acid, Gadoxetate disodium, Primovist®, Eovist®, Bayer HealthCare Pharmaceuticals, Leverkusen, Germany) remained available for liver imaging despite its linearity because it “met an important diagnostic need” without which, differential diagnosis could not be ascertained.^[12]^

MBCA alternatives for GBCAs, including Gd(III)-EOB-DTPA, have previously been explored, but these typically consist of Mn(II), *q* = 1 complexes that exhibit lower relaxivities relative to their GBCA counterparts, thereby requiring administration of higher doses to generate sufficient contrast enhancement.^[13]^ For instance, mangafodipir trisodium (MnDPDP) was designed as a gadolinium-free substitute for liver imaging but required two to four multiples (0.05-0.1 mmol/kg) of an already large dose of Gd(III)-EOB-DTPA (0.025 mmol/kg) to provide sufficient liver contrast in patients.^[14]^ Notwithstanding, optimization of the chemical chelator and structure chelating the paramagnetic center can improve relaxivity to generate next-generation Gd(III)-free contrast agents that outperform their GBCA counterparts.

More specifically, Gd(III)-EOB-DTPA generates targeted contrast in the liver through uptake by organic anion-transporting polypeptide 1 (Oatp1) proteins that are expressed on the membranes of human hepatocytes, namely *Oatp1b1* and *Oatp1b3*.^[15]^ The absence of these proteins, indicated by low enhancement following Gd(III)-EOB-DTPA administration, thereby allows for detection and characterization of metastases in the liver, nodular lesions of cirrhosis, as well as quantitative assessment of liver perfusion and hepatocyte function in diffuse liver diseases.^[16]^ In the preclinical context, Gd(III)-EOB-DTPA provides liver contrast through uptake via the rat and murine ortholog transporter, named *Oatp1a1* for both species.^[17]^ Beyond endogenous expression for liver imaging, this protein-ligand pair has been developed as an MRI reporter gene system, although its Gd(III)-based probe is identified as a limitation to its clinical translation for *in vivo* tracking of gene- and cell-based therapies in patients.^[18]^

Here we sought to develop an Oatp1-targeted MBCA with high relaxivity. We based our contrast agent design on a porphyrin ligand, which consists of four pyrroles that altogether make up a large octahedral ring molecule, chelated to a high spin (*S* = 2) manganese(III) center (**Figure 1**). Previously, combining the electron-spin properties of Mn(III) paramagnetic label with the rigid, geometrically flat porphyrin chelator allowed for water coordination both above and below the plane of the molecule (*q* = 2), resulting in increased relaxivity without sacrificing thermodynamic stability^[19]^. Curiously, Mn(III)Ps also stray in behavior from GBCAs and other Mn(II) agents by exhibiting an increase in relaxivity above field strengths of 2 MHz instead of a steep decline,^[20]^ further compelling us to employ this structure for Oatp1 imaging. To address the need for an Oatp1-specific Gd(III)-free contrast agent without compromising relaxation efficiency, we designed and synthesized Mn(III) 20-(4-ethoxyphenyl)porphyrin-5,10,15-tricarboxylate (Mn(III)TriCP-PhOEt). We report its superiority to Gd(III)-EOB-DTPA with respect to relaxivity at clinical field strengths, demonstrate its specificity as an MR probe for *Oatp1a1* and *Oatp1b3* transporters, as well as its effectiveness for *in vivo* contrast-enhanced liver imaging in mice.

**Figure 1.**
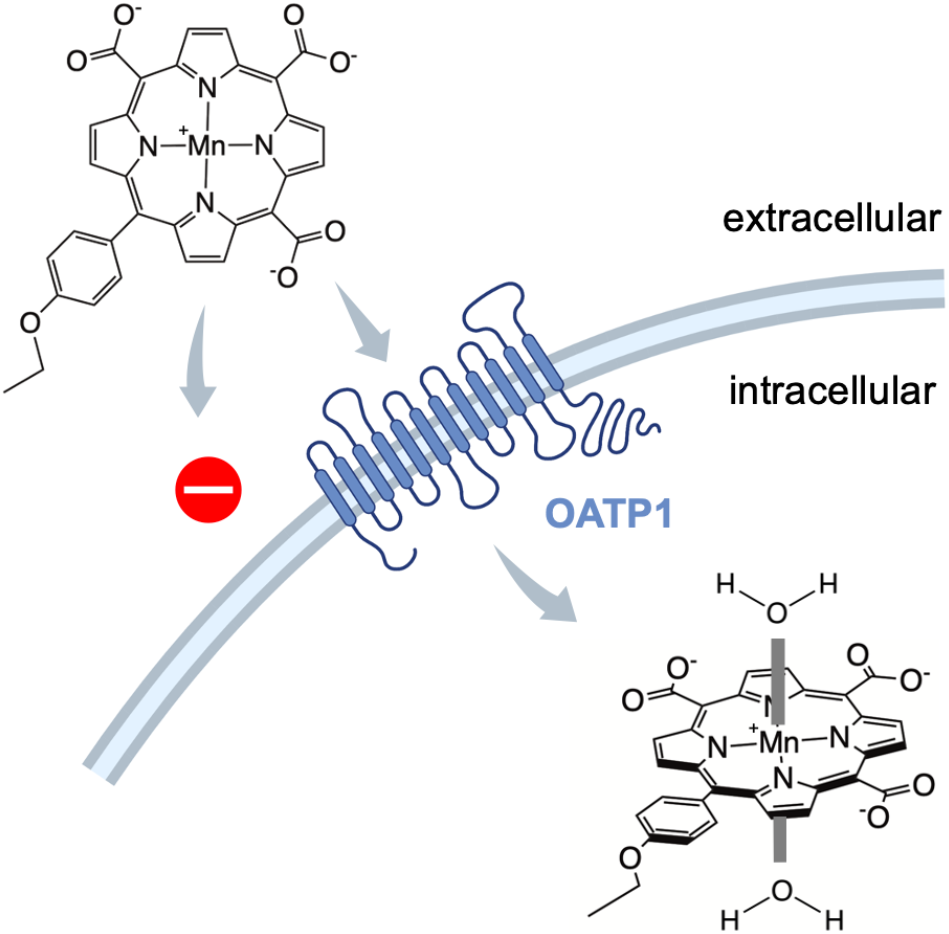
Oatp1-mediated Transport of Mn(III)TriCP-PhOEt for Targeted Magnetic Resonance Imaging. Cellular membranes are impermeable to Mn(III)TriCP-PhOEt. However, Oatp1a1 (rat) and/or Oatp1b3 (human) transporters that are natively expressed on hepatocytes recognize Mn(III)TriCP-PhOEt as a ligand and transition the contrast agent to the cellular cytoplasm. The planar geometry of Mn(III)TriCP-PhOEt allows for water coordination both above and below the paramagnetic center. Its accumulation in targeted cells would result in significant shortening of the longitudinal relaxation time (*T*_1_), thereby allowing Oatp1a1 and/or Oatp1b3-expressing cells to appear bright on *T*_1_-weighted magnetic resonance images. Molecules are not drawn to scale.

## RESULTS

### Synthesis of Mn(III)TriCP-PhOEt

Manganese (III) 20-(4-ethoxyphenyl)porphyrin-5,10,15-tricarboxylate (Mn(III)TriCP-PhOEt) was synthesized using an acid catalyzed condensation reaction between aldehydes and pyrrole (**Figure 2A**). Complete details of synthesis procedures and instrumentation are located in *Supporting Information*. All generated compounds were validated with ESI-MS and/or ^1^H NMR (**Supplementary Figures 1–4**). The synthetic precursors, α-1H-pyrrol-2-yl-1H-pyrrole-2-acetic acid ethyl ester (EDP) and 2,2′-(4-ethoxyphenylmethylene)bis-1H-pyrrole (DPM-PhOEt) were produced at 79% and 67.9% yield, respectively. Coupling of EDP and DPM-PhOEt gave triethyl 20-(4-ethoxyphenyl)porphyrin-5,10,15-tricarboxylate (TriEP-PhOEt) at 23% yield (484.5 mg). Upon metal-insertion, manganese (III) triethyl 20-(4-ethoxyphenyl)porphyrin-5,10,15-tricarboxylate (MnTriEP-PhOEt) was produced at 46.6% yield which was then hydrolyzed to generate final product, Mn(III)TriCP-PhOEt at 82.7% yield. ESI-MS of Mn(III)TriCP-PhOEt found a mass-to-charge ratio (*m/z*) of 306.1 ([M]^2−^), in close agreement with the *m/z* calculated for C_31_H_17_MnN_4_O_7_^2−^of 306.03 (**Figure 2B**). TriEP-PhOEt, Mn(III)TriEP-PhOEt, and Mn(III)TriCP-PhOEt exhibited unique peaks on UV-Vis spectra at 414 nm, 460 nm, and 467 nm, respectively (**Figure 2C**). Dominant peak of Mn(III)TriCP-PhOEt on HPLC-UV exhibited 97.8% purity (**Figure 2D**).

**Figure 2.**
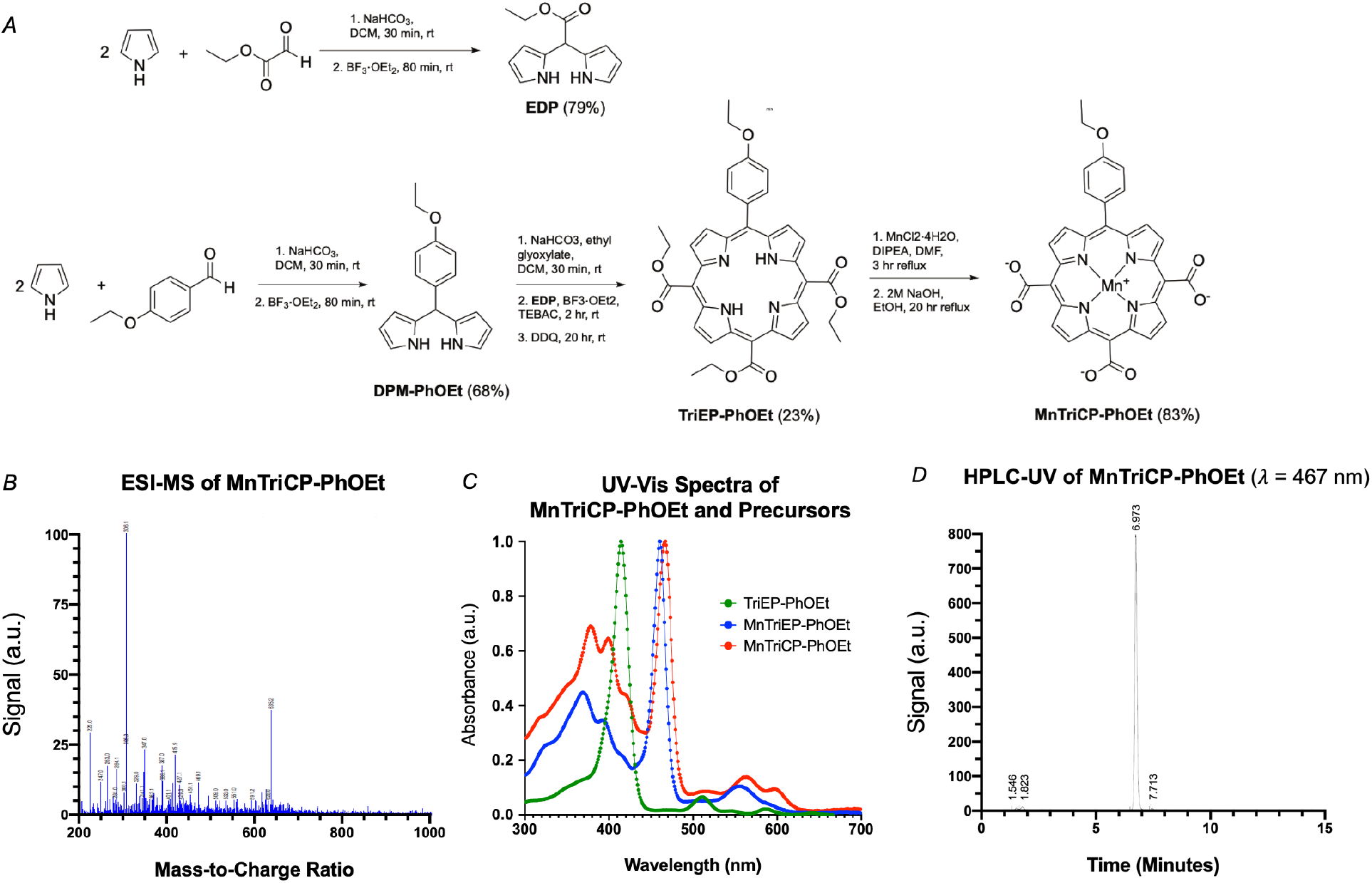
Synthesis and Validation of Mn(III)TriCP-PhOEt. A, Synthesis scheme for Manganese (III) 20-(4-ethoxyphenyl) porphyrin-5,10,15-tricarboxylate (MnTriCP-PhOEt). α-1H-Pyrrol-2-yl-1H-pyrrole-2-acetic acid ethyl ester, EDP. 2,2′-(4-Ethoxyphenylmethylene) bis-1H-pyrrole, DPM-PhOEt. Triethyl 20-(4-ethoxyphenyl) porphyrin-5,10,15-tricarboxylate, TriEP-PhOEt. B, Electrospray ionization mass spectrometry (ESI-MS) of Mn(III)TriCP-PhOEt, negative mode. Observed *m/z* = 306.1 ([M]^2−^). Calculated *m/z* for C_31_H_17_MnN_4_O_7_^2−^= 306.03. C, UV spectra of TriEP-PhOEt 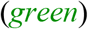, Mn(III)TriEP-PhOEt 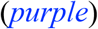, and Mn(III)TriCP-PhOEt 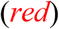. D, HPLC-UV of Mn(III)TriCP-PhOEt, detected at 467 nm.

### Mn(III)TriCP-PhOEt exhibits increased *r*_1_ relaxivity relative to Gd(III)-based contrast agents

Nuclear magnetic relaxation dispersion (NMRD) profiles were acquired at *B*_0_ ∈ (0, 1) T field strengths, 1 mM concentrations, and 37°C temperatures using a Spinmaster fast field-cycling relaxometer (Stelar s.r.l, Mede Italy). The relaxivities of Gd(III)-DTPA and Gd(III)-EOB-DTPA acquired were in agreement with literature values.^[21]^ At low fields, where *B*_0_ < 0.01 T, all three contrast agents exhibited linear *r*_1_ relaxivities with respect to variable field strength. For *B*_0_ ∈ (0, 0,01) T, the mean average *r*_1_ relaxivity for Gd(III)-DTPA was 7.817 ± 0.068 mmol^−1^ s^−1^, 9.955 ± 0.087 mmol^−1^ s^−1^ for Gd(III)-EOB-DTPA, and 5.911 ± 0.037 mmol^−1^ s^−1^ for Mn(III)TriCP-PhOEt (**Figure 3A**). At 0.01 < *B*_0_ < 1 T, the *r*_1_ relaxivity of both GBCAs decreases to 4.763 ± 0.042 mmol^−1^ s^−1^ for Gd(III)-DTPA, and 7.330 ± 0.045 mmol^−1^ s^−1^ for Gd(III)-EOB-DTPA. However, across this same range, the *r*_1_ relaxivity of Mn(III)TriCP-PhOEt increases and surpasses that of Gd(III)-DTPA at 0.1 T and that of Gd(III)-EOB-DTPA at 0.18 T, reaching a maximum of 8.942 ± 0.091 mmol^−1^ s^−1^ at 0.425 T before decreasing marginally to 8.794 ± 0.071 mmol^−1^ s^−1^ at 1 T (**Figure 3A**). Inversion recovery experiments (n=3) were performed at 1.5 T on a clinical MRI system at 18°C to calculate relaxivities at those field strengths. At 1.5 T, Mn(III)TriCP-PhOEt displays a significantly higher relaxivity (8.32 ± 0.21 mmol^−1^ s^−1^) than both Gd(III)-DTPA (4.66 ± 0.34 mmol^−1^ s^−1^, 56.02% of *r*_1MnTriCP-PhOEt_, p<0.0001) and Gd(III)-EOB-DTPA (7.32 ± 0.36 mmol^−1^ s^−1^, 88.01% of *r*_1MnTriCP-PhOEt_, p=0.0178) (**Figure 3A**). At 3 T, Mn(III)TriCP-PhOEt also displays a significantly higher relaxivity (11.25 ± 0.35 mmol^−1^ s^−1^) than Gd(III)-DTPA (6.20 ± 0.18 mmol^−1^ s^−1^, 55.11% of *r*_1MnTriCP-PhOEt_, p<0.0001) and Gd(III)-EOB-DTPA (8.34 ± 0.23 mmol^−1^ s^−1^, 74.13% of *r*_1MnTriCP-PhOEt_, p<0.0001) (**Figure 3B**).

**Figure 3.**
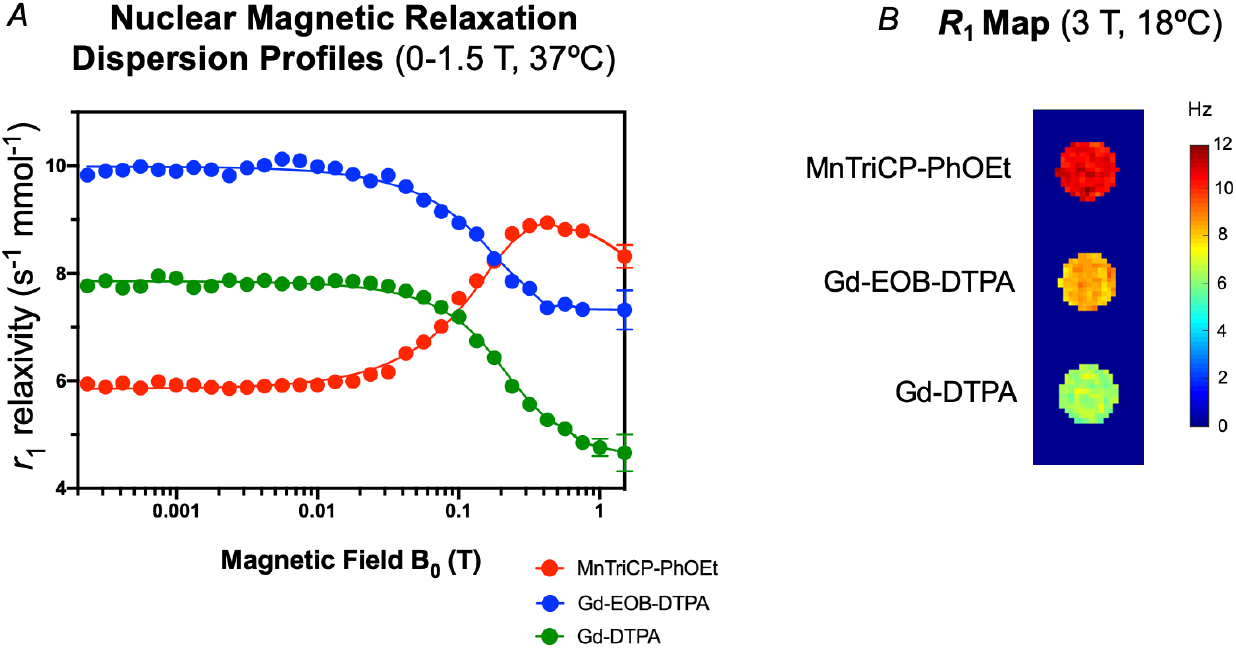
Paramagnetic Characterization of Mn(III)TriCP-PhOEt at Various Field Strengths. A, Nuclear magnetic relaxation dispersion profiles of 1 mM solutions of Mn(III)TriCP-PhOEt 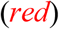, Gd(III)-EOB-DTPA 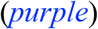, and Gd(III)-DTPA 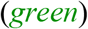 at *B*_0_ ∈ (0, 1.5) T, 37°C. Error bars indicate standard deviation. Cubic spline regressions are plotted for each contrast agent. B, *R*_1_ relaxation (Hz) map of 1 mM solutions of MnTriCP-PhOEt, Gd(III)-EOB-DTPA, and Gd(III)-DTPA at *B*_0_ = 3 T, 18°C.

### Cells expressing Oatp1a1/1b3 transporters exhibit increased *r*_1_ relaxivity when treated with Mn(III)TriCP-PhOEt

Rat C6 glioma cells and human triple negative breast cancer MDA-MB-231 cells were engineered with lentivirus to express Oatp1a1 with tdTomato and Oatp1b3 with zsGreen, respectively (**Figure 4A, 4B**). Cells were sorted with fluorescent activated cell sorting (FACS) resulting in an engineered cell population with >99% purity (**Figure 4C, 4D**). Control and transporter-expressing cells were treated in suspension with Mn(III)TriCP-PhOEt, Gd(III)-EOB-DTPA, or Gd(III)-DTPA, subsequently washed, trypsinized and pelleted for *R*_1_ measurements at 3 T (n=3). As a control, naive and transporter-expressing cells did not exhibit differences in spin-lattice relaxation rates (Hz) when incubated with 1-mM Gd(III)-DTPA, which is not permeable to the cell membrane, and is not recognized as a ligand by either Oatp1a1 or Oatp1b3 transporters (**Figure 4E**, *first column*). Additionally, control cells not expressing Oatp1a1 or Oatp1b3 transporters did not exhibit differences in relaxivity when treated with either Gd(III)-DTPA, Gd(III)-EOB-DTPA, or Mn(III)TriCP-PhOEt, confirming that all three contrast agents are impermeable to cell membranes in the absence of Oatp1a1 or Oatp1b3 transporters (**Figure 4E**, *first row*). For the rat Oatp1a1 transporter, treatment with Mn(III)TriCP-PhOEt resulted in an increased spin-lattice relaxation rate (2.76 ± 0.38 Hz) relative to Oatp1a1-expressing cells treated with Gd(III)-EOB-DTPA (2.50 ± 0.36 Hz), albeit this difference was not statistically significant (p=0.845) (**Figure 4E, 4F**). For the human Oatp1b3 transporter, cells treated with Mn(III)TriCP-PhOEt exhibited a significantly increased spin-lattice relaxation rate (2.24 ± 0.13 Hz) relative to the same cells treated with Gd(III)-EOB-DTPA (1.52 ± 0.03 Hz, p=0.0429) (**Figure 4E, 4F**).

**Figure 4.**
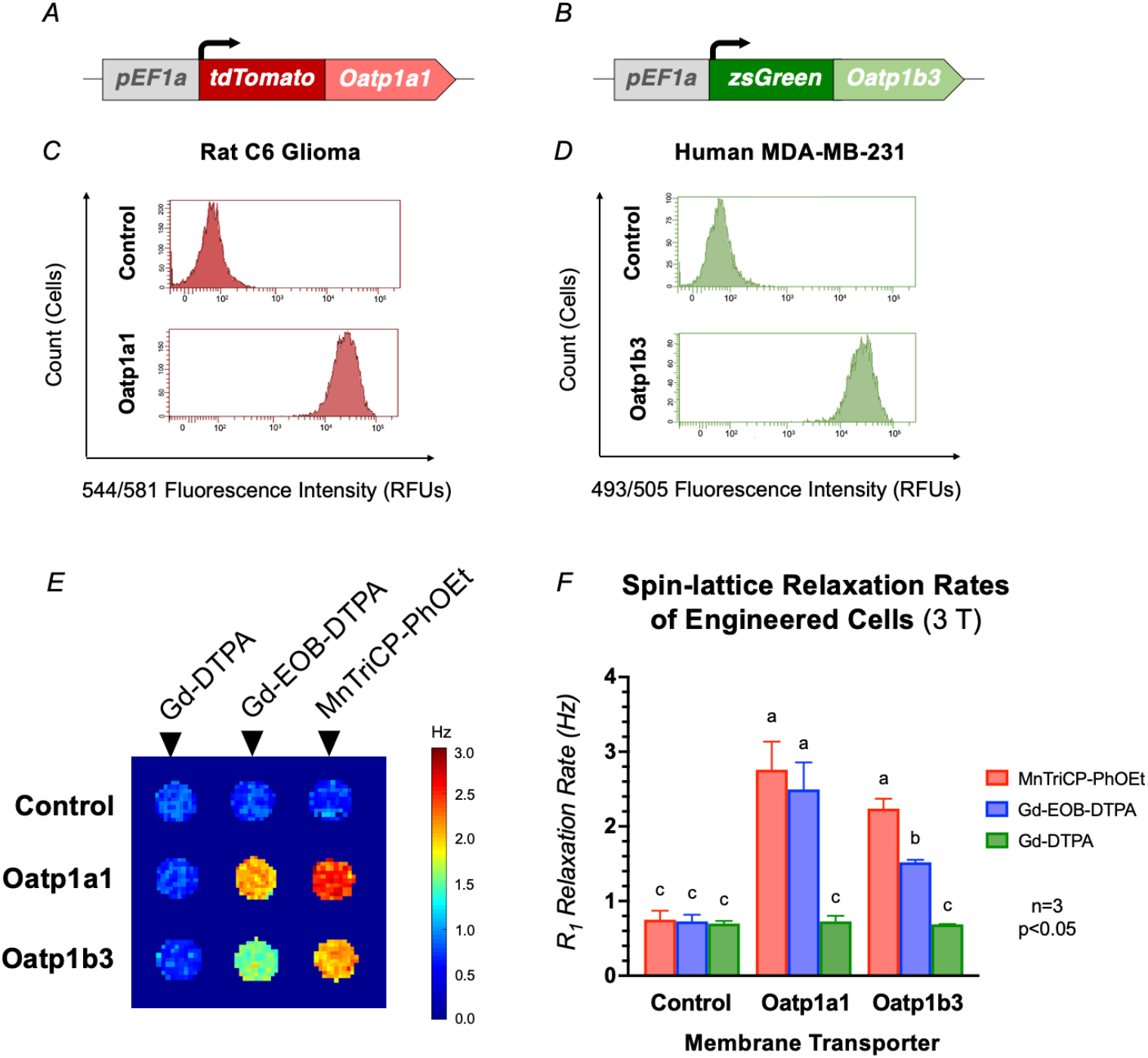
Mn(III)TriCP-PhOEt Increases in Spin-Lattice Relaxation Rates of Cells Expressing Oatp1a1 and Oatp1b3 Liver Transporters. C6 rat glioma cells were engineered to express the rat liver transporter Oatp1a1 with tdTomato (A) and human breast cancer cells (MDA-MB-231) were engineered to express the human liver transporter Oatp1b3 with zsGreen (B). Flow cytometry of control and engineered rat cells (C) and human cells (D) exhibiting pure populations of transporter-expressing cells. RFU, relative fluorescent units. Maps of spin-lattice relaxation rates (Hz) for control, Oatp1a1-expressing, and Oatp1b3-expressing cells incubated with either Gd(III)-DTPA, Gd(III)-EOB-DTPA, or Mn(III)TriCP-PhOEt for 1 hour were acquired at 3 Tesla (E). Average spin-lattice relaxation rates (Hz) of cells treated with contrast agents (n=3) were computed from relaxation rate maps (F). Error bars indicate one standard deviation.

### Mn(III)TriCP-PhOEt enables *in vivo* contrast enhancement of liver tissue at clinical doses

As a control, manganese(III) porphyrin-5,10,15,20-tetracarboxylate (MnTCP), which is structurally related to Mn(III)TriCP-PhOEt barring the presence of a fourth carboxylic acid functional group in place of the PhOEt moiety, exhibited significant contrast enhancement in the kidneys, but virtually no *T*_1_-shortening effects in liver tissue (**Supplementary Figure 5**) analogous to Gd(III)-DTPA, and in agreement with previous Mn(III)TCP biodistribution findings.^[22]^ Upon intraperitoneal injection of 0.025 mmol/kg Mn(III)TriCP-PhOEt, contrast enhancement was observed for *T*_1_-weighted images of the liver and gallbladder as part of the hepatobiliary system and the kidneys and bladder as part of the renal system (**Figure 5A**). On average across all mice (n=4), a significant increase in liver signal intensity (arbitrary units, a.u.) was observed between pre-contrast images and post-contrast images acquired greater than 28 minutes following intraperitoneal injection of 0.025 mmol/kg Mn(III)TriCP-PhOEt (171.2 ± 39.6%, p=0.031), which continued to increase until a maximal increase in liver signal intensity (a.u.) of 191.4 ± 37.3% was observed at the 52-minute imaging timepoint (p=0.014) (**Figure 5B**). The average area under the curve (AUC) within a 60-minute imaging course (n=3) between the liver (29,298 ± 1639 a.u.) and kidney volumes (30,442 ± 758 a.u.) were not significantly different (p=0.607), but both were significantly higher than the average AUC of the heart (12,087 ± 1642 a.u., p<0.0001) (**Figure 5C, 5D**). The AUC_liver_ to AUC_kidney_ ratio of 1.00 to 1.04 suggests that the percentage of Mn(III)TriCP-PhOEt clearance by organ is relatively equal between the hepatobiliary and renal systems.

**Figure 5.**
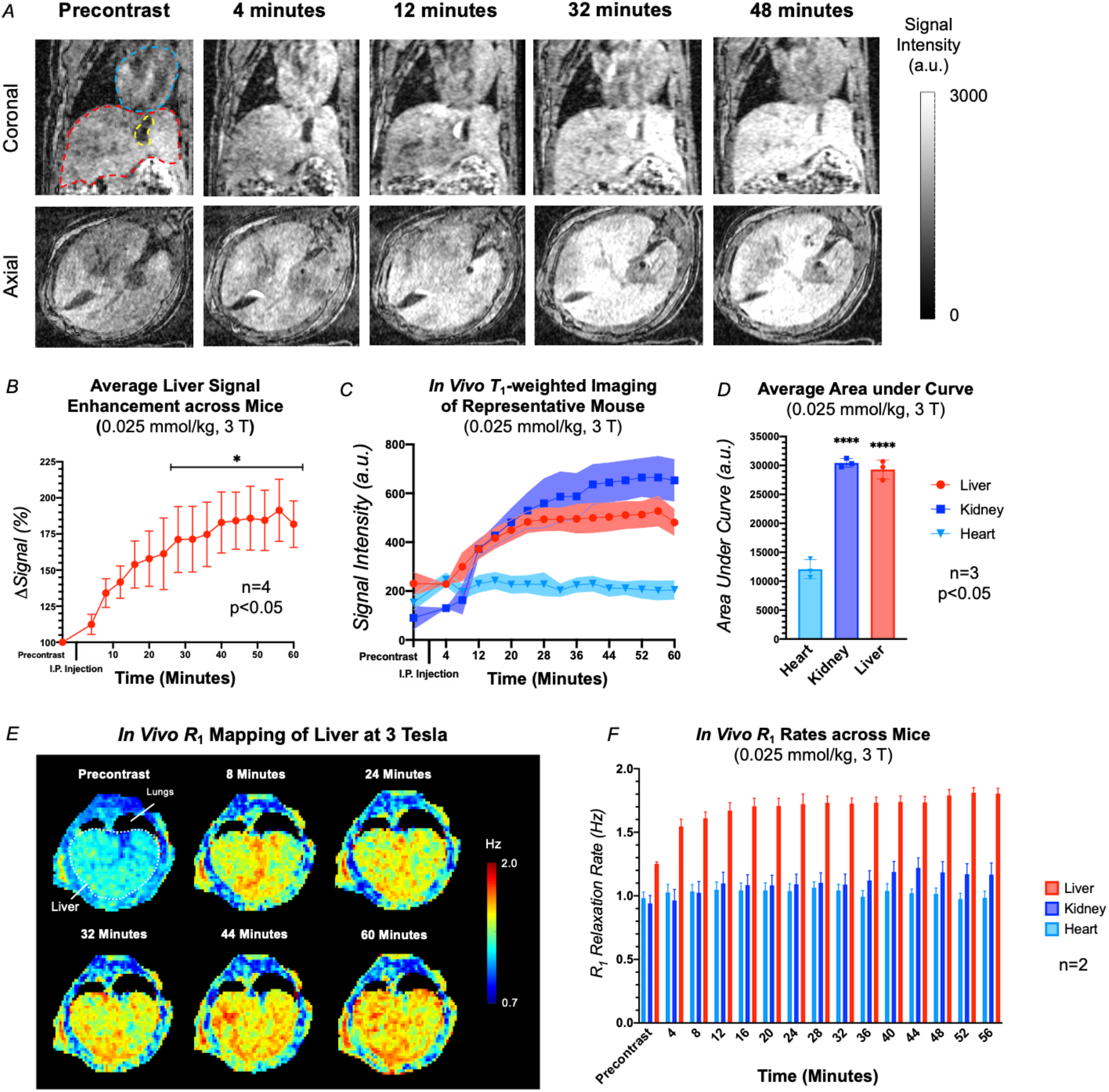
*T*_1_-weighted and Quantitative Liver Imaging in Mice with 0.025 mmol/kg Mn(III)TriCP-PhOEt at 3 Tesla. A, *T*_1_-weighted images of representative mouse before and following injection of 0.025 mmol/kg Mn(III)TriCP-PhOEt. B, Signal intensity, arbitrary units (a.u.). C, Dynamic signal intensity curves of liver 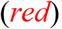, kidney 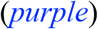, and heart 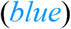 tissue of representative mouse before and following injection of 0.025 mmol/kg Mn(III)TriCP-PhOEt. Error bars represent standard deviation. D, Average change in signal intensity of liver tissue following injection of 0.025 mmol/kg Mn(III)TriCP-PhOEt, in reference to pre contrast value (n=4, ****p<0.0001). E, Representative *in vivo R*_1_ map of axial cross-section of mouse before and following injection of 0.025 mmol/kg Mn(III)TriCP-PhOEt, measured in hertz (Hz). F, Average *in vivo* longitudinal relaxation rate (Hz) of various tissues before and following intraperitoneal injection of 0.025 mmol/kg Mn(III)TriCP-PhOEt. Error bars represent standard error of the mean (n=2).

### Quantitative *R*_1_ mapping increases liver imaging sensitivity and specificity of Mn(III)TriCP-PhOEt relative to *T*_1_-weighted imaging

In addition to dynamic contrast enhanced *T*_1_-weighted imaging, dynamic contrast enhanced quantitative imaging was performed to measure *in vivo* changes in spin-lattice relaxation rates before and up to an hour following 0.025 mmol/kg Mn(III)TriCP-PhOEt administration (**Figure 5E**). Whereas significant differences in liver signal intensity on *T*_1_-weighted images were observed 28 minutes after administration, significantly lower *T*_1_ values of liver tissue was observed as early as 12 minutes post administration (598.5 ± 52.9 ms, p=0.012) relative to the *T*_1_ time of the liver precontrast (798.5 ± 16.9 ms) (**Figure 5F**). Moreover, the average percent standard deviation of liver signal intensity on *T*_1_-weighted images (19.02 ± 4.3%) was significantly greater than the average percent standard deviation of liver spin-lattice relaxation rates observed on *R*_1_ maps (7.06 ± 2.1%, p=0.010). This increased precision in the latter method enables more sensitive detection of changes relative to pre-contrast images. Slight decreases in *T*_1_ times were observed for both kidney and heart tissue but no significant differences were observed for either tissue at any timepoint during the 60-minute course of imaging, relative to precontrast values, even after segmenting each organ into independent compartments *e.g.* renal medulla, renal cortex, heart walls, heart chambers.

### *Ex vivo* nuclear magnetic relaxation dispersion confirms spin-lattice relaxation rate differences are liver-specific

Mice (n=3) were intraperitoneally injected with either saline or 0.025-mmol/kg Mn(III)TriCP-PhOEt and sacrificed 36 minutes post administration. Liver, kidney and muscle tissue was harvested and NMRD was performed for each tissue sample. Significant differences in spin-lattice relaxation times were observed specifically between liver tissue of Mn(III)TriCP-PhOEt and control mice at 0.3 < *B*_0_ < 1 T (p<0.05, **Figure 6A**), which is not unexpected given that the relaxivity of Mn(III)TriCP-PhOEt decreases drastically at field strengths lower than this range (**Figure 3A**). Following the same finding observed *in vivo* at 3 T however (**Figure 5F**), no significant differences were observed at 0.01 < *B*_0_ < 1 T for either kidney or muscle tissue between Mn(III)TriCP-PhOEt and control tissue samples (**Figure 3A**). The maximal difference in spin-lattice relaxation rate was 0.679 Hz for liver tissue at *B*_0_ = 0.3057 T, 0.452 Hz for kidney tissue at *B*_0_ = 0.0154 T, and 0.297 Hz for muscle tissue at *B*_0_ = 0.0154 T.

**Figure 6.**
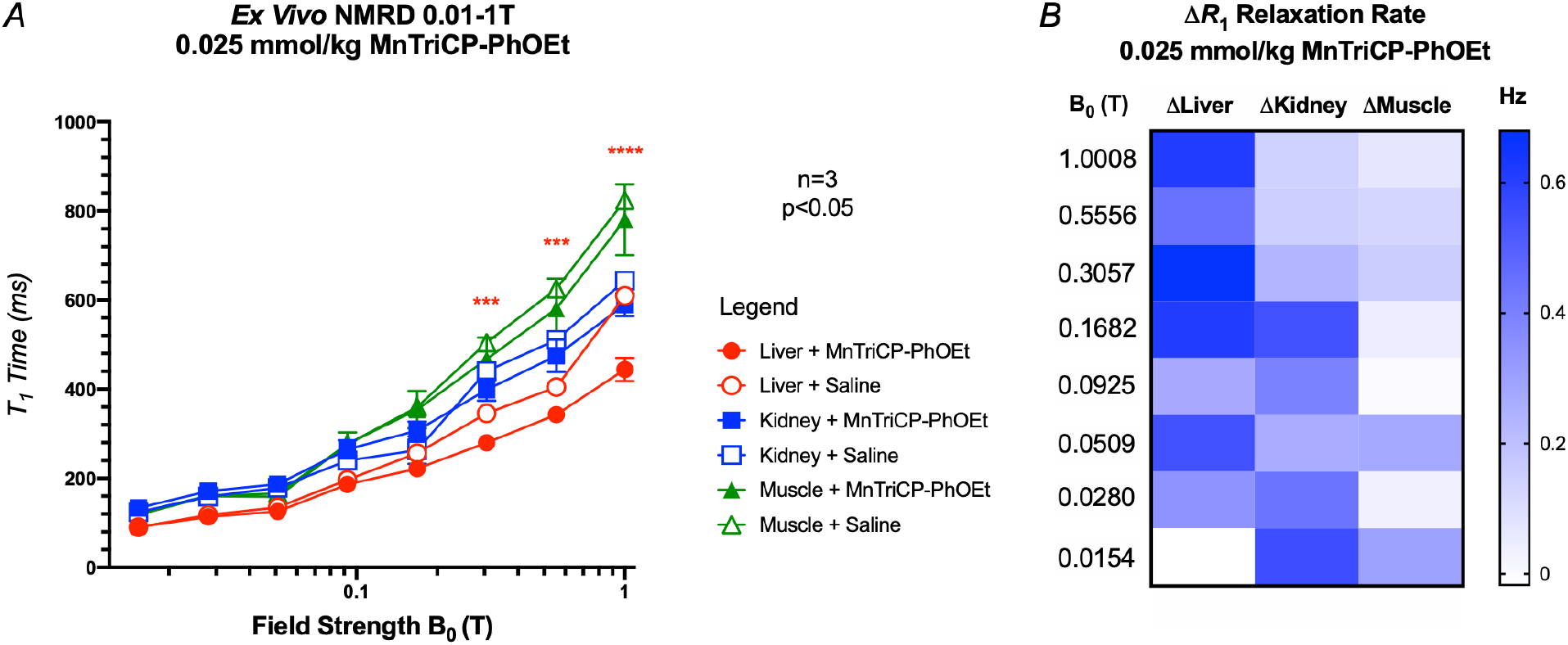
*Ex Vivo* Nuclear Magnetic Relaxation Dispersion of Various Tissues following Mn(III)TriCP-PhOEt Administration. A, Nuclear magnetic relaxation dispersion (NMRD) profiles of tissues from mice (n=3) collected 36 minutes following intraperitoneal injection of 0.025 mmol/kg Mn(III)TriCP-PhOEt or saline (control) acquired at low field. B, Heat map of differences in spin-lattice relaxation rates (Hz) between mice administered Mn(III)TriCP-PhOEt or saline.

## DISCUSSION

In response to increased evidence of long-term tissue retention, a new class warning for all GBCAs was announced by the FDA in 2017^[11]^ and the European Medicines Agency (EMA) followed by suspending marketing authorizations for three of the five GBCAs comprised of linear chelators due to their increased susceptibility to release Gd(III).^[12]^ However, Gd(III)-EOB-DTPA remained available for liver imaging because it provided unique diagnostic information that could not be obtained by alternative methods.^[12]^ Here, we demonstrate that Mn(III) 20-(4-ethoxyphenyl)porphyrin-5,10,15-tricarboxylate (Mn(III)TriCP-PhOEt) offers high specificity and high sensitivity for *Oatp1*-targeted imaging *in vivo*, which enables contrast-enhanced liver imaging on MR^[15]^ and paves the path towards *Oatp1* reporter gene imaging^[18a, 18c, 23]^ without the need to include Gd(III) in the protocol or sacrifice relaxivity (**Figure 1**).

First, the Mn(III)TriCP-PhOEt complex was shown to have superior *r*_1_ relaxivity at 1.5 Tesla, and even more so, at 3 Tesla, relative to that of Gd(III)-EOB-DTPA (**Figure 3**). Next, we demonstrated the specificity of its uptake through the rodent *Oatp1a1* and human *Oatp1b3* transporters using cells genetically engineered to synthetically express these proteins. The Mn(III)TriCP-PhOEt complex again outperformed the contemporary Gd(III)-EOB-DTPA probe, with respect to resultant *r*_1_ relaxivities of Oatp1-expressing cells incubated with each respective agent (**Figure 4**). Finally, uptake of Mn(III)TriCP-PhOEt *in vivo* by murine liver cells that endogenously express *Oatp1a1* was demonstrated via significant increases in liver signal intensity following low-dose intraperitoneal injections (0.025 mmol/kg) of Mn(III)TriCP-PhOEt (**Figure 5A-D**).

Dynamic *R*_1_ mapping at 3 Tesla was additionally performed to quantify *T*_1_-shortening effects following Mn(III)TriCP-PhOEt administration, and *T*_1_-shortening in the liver was significantly more pronounced relative to the kidneys (**Figure 5E, 5F**). Although this was in discordance with the almost equivalent increases in signal intensity observed in the liver and kidneys on *T*_1_-weighted images, this finding was further confirmed by *ex vivo* NMRD measurements (**Figure 6**) and is likely due to the considerable variation in *T*_1_ values within the kidney volume as a reflection of its complex structure,^[24]^ and despite efforts to delineate individual renal compartments, the large variation in kidney *T*_1_ time persisted. Altogether, these findings support Mn(III)TriCP-PhOEt as an effective contrast agent for liver imaging on MR.

One immediate point of improvement, however, will be to increase the relatively low aqueous solubility of the complex, which was found to saturate at about 1.6 mM concentrations in saline at 37°C. Despite basing the design of this Mn(III)P on a less established anionic parent with carboxylic acid moieties at its periphery,^[22]^ with the notion that solubility would not present as an issue in animal studies, the overall aromatic structure of the porphyrin as well as the addition of the *Oatp1a1/1b3*-specific phenyloxy ethyl ligand contributed sufficiently to its hydrophobicity. This was mitigated to some extent by dissolving Mn(III)TriCP-PhOEt in dimethyl sulfoxide (DMSO) before further dilution in saline prior to injection.

Yet even when dissolved in organic solvents, intravenous injections were not feasible considering the volume of injection required, thereby warranting intraperitoneal administrations in our animal studies, which non-negligibly reduced the bioavailability of the contrast agent in the blood.^[25]^ Despite this, significant increases in the mean signal intensity and *R*_1_ rates of the liver volume were observed in mice, probably owed to the high *r*_1_ relaxivity of Mn(III)TriCP-PhOEt. We anticipate that improving solubility to enable intravenous injections in animals would further amplify contrast enhancement, though the human equivalent dose, which is typically more than 10-fold smaller than that of rodent animal models,^[26]^ may be small enough to already permit intravenous injections in patients without the need to increase solubility.

Overall, Mn(III)Ps offer a unique combination of advantages as MRI contrast agents. Mn(III)Ps have been established as thermodynamically stable and kinetically inert, but even so, manganese, unlike Gd(III), is a nutritional element that the human body is able to physiologically incorporate or excrete, thereby evading concerns of long-term accumulation and subsequent toxicity.^[27]^ In extreme cases, Mn ion exposure has been associated with a neurodegenerative disorder called manganism, but a large cumulative amount of free Mn (over 3.8 mmol/kg) is required to induce toxicity, unlikely to be achieved even after repeat Mn(III)P doses.^[28]^ In fact, manganese enhanced MRI (MEMRI) studies, whereby small doses of free Mn are administered directly to preclinical animals, is routinely used to map neural activity.^[29]^ Even in humans, Mn(II)-DPDP by design operated on the mechanism of Mn ion release, and was clinically assessed as having a safety factor (LD_50_/effective dose) of 540, significantly greater than that of GBCAs that have safety factors ranging from 60 to 100.^[30]^ For our own work, future studies will focus on developing a more soluble *Oatp1*-targeted Mn(III)P and characterizing its kinetic and thermodynamic stability as well as assessing its performance in detecting liver metastasis in mice and engineered cells for reporter gene tracking.

Mn(III)Ps have been extensively studied *in vitro*^[19]^ though the precise mechanics dictating their “anomalously” high relaxivity are less deeply explored than those of Gd(III) and Mn(II), owing to the unusual electron spin behaviors and strong metal-ligand orbital interactions.^[20, 31]^ Early *in vivo* imaging studies have focused on Mn(III)Ps as vascular agents, and more recently, works have expanded functions of Mn(III)Ps for extracellular, intracellular and intravascular applications.^[27, 32]^ The next generation of contrast agents will have high tissue specificity and/or the ability to act as reporters of the environment in which they are distributed. With four distinct modification sites at the porphyrin *meso*-positions, Mn(III)Ps will be exceptionally suited to facilitate these efforts. In fact, oxygen,^[33]^ zinc,^[19]^ calcium,^[34]^ secreted alkaline phosphatase,^[35]^ and esterase-responsive^[36]^ Mn(III)Ps have already been reported. Historically, GBCAs have shown tremendous utility and have single-handedly enabled contrast enhanced MR imaging in patients for many years, but as more insight into their retention in patients’ bodies and the environment is revealed, their continued use loses justification, especially as contrast agents that are both safer and better-performing arise.

## CONCLUSION

Our study finds that Mn(III) 20-(4-ethoxyphenyl)porphyrin-5,10,15-tricarboxylate (MnTriCP-PhOEt) exhibits superior *r*_1_ relaxivity relative to Gd(III)-EOB-DTPA at clinically-relevant field strengths and demonstrates specificity to rodent *Oatp1a1* and human *Oatp1b3* channels as biomarkers of hepatocyte viability. In mice, 0.025 mmol/kg injections of Mn(III)TriCP-PhOEt enhance liver contrast with high sensitivity and specificity in 3-dimensions. Mn(III)TriCP-PhOEt offers a new approach for liver imaging on MRI without the need to administer gadolinium to patients or sacrifice sensitivity.

## ACKNOWLEDGEMENTS

Financial support for this manuscript was provided by Natural Sciences and Engineering Research Council (NSERC) Discovery Grants (J.A.R. RGPIN-2016-05420, T.J.S. RGPIN-2017-06338, and X.A.Z. RGPIN-2016-489075), Ontario Institute for Cancer Research Investigator Award (T.J.S. IA-028) and Canadian Institutes of Health Research Project Grant (J.A.R. 377071). X.A.Z. is also grateful to the University of Toronto Scarborough, Canada Foundation for Innovation, and Ontario Research Fund for their support. This work was additionally supported by the Breast Cancer Society of Canada (N.N.N.), and an NSERC Postgraduate Doctoral Scholarship (N.N.N.).

## SUPPORTING INFORMATION

### Chemical Reagents and Instrumentation

All reagents and solvents were of commercial reagent grade and were used without further purification except where noted. Pyrrole, 4-ethoxybenzaldehyde, 4-methoxybenzaldehyde, BF_3_•OEt_2_, TEBAC, MnCl_2_•4H_2_O and DCM were obtained from Sigma Aldrich (St. Louis, Missouri, USA). DDQ was obtained from AK Scientific, Inc. (Union City, California, USA). Ethyl glyoxylate was obtained from Alfa Aesar (Haverhill, Massachusetts, USA). NaHCO_3_ was obtained from Caledon Laboratories, Ltd. (Georgetown, Ontario, Canada). DIPEA was purchased from Acros Organics (Fair Lawn, New Jersey, USA). HEPES buffer (25 mM, pH 7.4) fixed at the ionic strength of 0.1 M using NaCl was used for all measurements. All other reagents and solvents were purchased from Fischer Scientific Inc. (Hampton, New Hampshire, USA). Thin layer chromatography (TLC) was performed using TLC silica gel 60 F254 plates from Merck Group (Darmstadt, Germany). Column chromatography was set up using Silica Gel 60, 250-400 microns from Desican Inc. (Scarborough, Ontario, Canada). Cation ion exchange was performed using an Amberlite® IR120, H resin (Sigma Aldrich). Dialysis was performed by use of Spectra/Por® 3 Dialysis membrane 1K MWCO (Fischer Scientific Inc.).

### α-1H-Pyrrol-2-yl-1H-pyrrole-2-acetic acid, ethyl ester (EDP)

The reaction was carried out in the dark under room temperature. 10 g (120 mmol) NaHCO_3_ was mixed with 100 mL DCM under argon for 20 minutes. 20 mL (287 mmol) distilled pyrrole and 4 mL (20.2 mmol) ethyl glyoxylate (50% v/v in toluene) was then added and stirred for 30 minutes. After TLC confirmation of full formation of intermediate alcohol product, the mixture was vacuum filtered to remove NaHCO_3_. 35 μL BF_3_•OEt_2_ was then added along with 100 mL DCM and the mixture stirred for another 60 minutes. The crude product was washed with 100 mL saturated NaHCO_3_ and dried over Na_2_SO_4_. The solvent was removed under reduced pressure. The product was purified by column chromatography (silica gel, elution solvent = 7:3 DCM/Hexanes). 3.50 g (79.4%) of **EDP** as a light pink solid was isolated. ^1^H-NMR (500 MHz, CDCl3): δ= 8.46 (s, 2H, -NH), 6.72 (q, 2H, pyrrole-αH), 6.16 and 6.08 (q, 4H, pyrrole-βHs), 5.5 (s, 1H, -CH), 4.23 (q, 2H, -CH_2_) and 1.31 (t, 3H,-CH_3_).

### 2,2′-(4-Ethoxyphenylmethylene)bis-1H-pyrrole (DPM-PhOEt)

The reaction was carried out in the dark under room temperature. 0.67 mL (4.8 mmol) 4-ethoxybenzaldehyde was mixed with 2.43 g (28.9 mmol) NaHCO_3_ in 25 mL DCM for 20 minutes under argon atmosphere. 5 mL (72 mmol) distilled pyrrole was then added and stirred for 30 minutes. After TLC confirmation of full formation of intermediate alcohol product, the mixture was vacuum filtered to remove NaHCO_3_. 32 μL BF_3_•OEt_2_ was then added along with 25 mL DCM and the mixture stirred for another 70 minutes. The crude product was washed with 100 mL saturated NaHCO_3_, and dried over Na_2_SO_4_. The solvent was removed under reduced pressure and remaining pyrrole removed by vacuum distillation. The product was purified by column chromatography (silica gel, elution solvent = 9:1 Hexanes/Acetone). 0.867 g (67.9%) of **DPM-PhOEt** as a yellow-pink solid was isolated. ^1^H-NMR (500 MHz, DMSO-d6): δ= 10.50 (s, 2H, -NH), 6.82-7.03 (dd, 4H, benzene H), 6.58 (m, 2H, pyrrole-αH), 5.88 (m, 2H, pyrrole-βH), 5.62 (m, 2H, pyrrole-βH), 5.27 (s, 1H, - CH), 3.97 (q, 2H, -OCH_2_-), 1.29 (t, 3H, -CH_3_).

### Triethyl 20-(4-ethoxyphenyl)porphyrin-5,10,15-tricarboxylate (TriEP-PhOEt)

The reaction was carried out in the dark at room temperature. 867.2 mg (3.3 mmol) **DPM-PhOEt** was mixed with 1.29 mL (6.5 mmol) ethyl glyoxylate, 821 mg (9.8 mmol) NaHCO_3_ and 4.4 mL DCM under argon atmosphere for 2 hours. The intermediate dicarbinol product was confirmed by TLC, and the NaHCO_3_ was removed via vacuum filtration. The dark red intermediate was mixed with 711.5 mg (3.3 mmol) **EDP** and 24 mg TEBAC in 1400 mL DCM and stirred under argon for 50 minutes, after which 65 μL BF_3_•OEt_2_ was then added. After an additional 50 minutes of stirring, 3.31 g (14.6 mmol) DDQ was then added, and the mixture was allowed to stir overnight. The dark purple-green crude was concentrated by rotary evaporation followed by filtration using a double-layer silica/alumina column, washed with 8:2 DCM/ethyl acetate. The filtered solution was then purified by column chromatography (silica gel, elution solvent = 99:1 DCM/ethyl acetate). The semi-purified product was then centrifuged in 8:2 hexanes/ethyl acetate and the pellet collected to afford 484.5 mg (23.0%) dark purple powder of **TriEP-PhOEt**. ^1^H-NMR (500 MHz, CDCl3): δ= 9.52 (s, 4H, pyrrole-βH), 9.38 (d, 2H, pyrrole-βH), 8.99 (d, 2H, pyrrole-βH), 8.09 (d, 2H, benzene H), 7.32 (m, 2H, benzene H), 5.07-5.13 (overlapping q, 6H, ester -OCH_2_-), 4.33-4.37 (q, 2H, ether -OCH_2_-), 1.78-1.84 (overlapping t, 9H, ester -CH_3_), 1.56-1.66 (t, 3H, ether -CH_3_), −3.06 (s, 2H, -NH).

### Manganese (III) triethyl 20-(4-ethoxyphenyl)porphyrin-5,10,15-tricarboxylate (MnTriEP-PhOEt)

198.5 mg (0.307 mmol) of **TriEP-PhOEt** was dissolved with 0.37 mL (2.2 mmol) DIPEA and 1.445 g (7.3 mmol) MnCl_2_•4H_2_O in 20 mL of DMF. The mixture was refluxed for 2 hours. The reaction was then stopped and allowed to stir in air for another 2 hours. DMF was removed under reduced pressure, and the crude product was purified by column chromatography (silica gel, elution solvent = 9.5:0.5 DCM/methanol) to afford 225.8 mg of **MnTriEP-PhOEt**. ESI-MS (positive mode, methanol) found m/z = 699.2 ([M]^+^); calculated for C_37_H_32_MnN_4_O_7_^+^ (m/z = 699.16).

### Manganese (III) 20-(4-ethoxyphenyl)porphyrin-5,10,15-tricarboxylate (MnTriCP-PhOEt)

225.8 mg (0.307 mmol) of **MnTriEP-PhOEt** was dissolved in 45 mL ethanol and 35 mL THF. A 75 mL solution of 2M NaOH was added to the mixture. The solution was refluxed for 24 hours with HPLC monitoring. The organic solvents were then removed under reduced pressure, and the crude product washed with ethyl acetate (5 x 100 mL). The aqueous mixture was acidified with 1 M HCl to pH 2.00 followed by centrifugation to remove the supernatant. Dialysis was carried out (MWCO 1K) on the pellet. Crude 5 was then passed through a cation exchange column (stationary phase: Amberlite® IR120 H resin, mobile phase: double distilled H_2_O) to remove excess Mn(II) salts, and the product lyophilized. 175.4 mg (82.7%) of brown powder was collected as **MnTriCP-PhOEt**. ESI-MS (negative mode, H_2_O) found m/z = 306.0 ([M]^2−^); calculated for C_31_H_17_MnN_4_O_72_^−^ (m/z = 306.02).

### Nuclear magnetic dispersion and magnetic resonance profiles

Contrast agents were first reconstituted in dimethyl sulfoxide (DMSO), warmed to 37°C and then diluted in Dulbecco’s Modified Eagle Media (DMEM, Wisent Inc., Saint-Jean-Baptiste, Quebec, Canada) supplemented with 10% fetal bovine serum (FBS) to a final working concentration of 1 mM. Spin-lattice relaxation times were acquired at magnetic fields from 230 μT to 1 T on a fast field-cycling NMR relaxometer (SpinMaster FFC2000 1T C/DC, Stelar, s.r.l., Mede, Italy) by changing the relaxation field in 30 steps, logarithmically distributed using an acquisition field of 380.5 mT, with temperature kept constant at 25°C. Data for each contrast agent was subsequently fit into a LOWLESS spline curve with 10 points in the smoothing window. To determine relaxation rates at higher field strengths, an inversion recovery experiment was performed on a 1.5-Tesla GE CVMR and a 3-Tesla GE MR750 clinical scanner (General Electric Healthcare, Milwaukee Wisconsin, USA) using a 32-channel phase array head coil (General Electric Healthcare, Milwaukee Wisconsin, USA). A Fast Spin Echo Inversion Recovery (FSE-IR) pulse sequence was used with the following parameters: matrix size = 256×256, repetition time (TR) = 5000 ms, echo time (TE) = 19.1 ms, echo train length = 4, number of excitations = 1, receiver bandwidth = 12.50 kHz, inversion times (TI) = 20, 35, 50, 100, 125, 150, 175, 200, 250, 350, 500, 750, 1000, 1500, 2000, 2500, 3000, in-plane resolution = 0.27 mm^2^, slice thickness = 2.0 mm. Spin-lattice relaxation rates were computed via MatLab (R2020a, MathWorks, Natick, Massachusetts, USA) by calculating the signal intensity pixelwise across the inversion time image series, followed by non-linear least-squares fitting of the data to the following equation to output the spin-lattice relaxation time (*T*_1_):

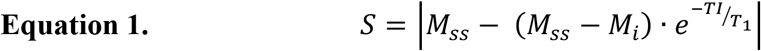

where *S*, *M*_*ss*_, and *M*_*i*_ represent the acquired signal, the longitudinal magnetization in steady state equilibrium, and the initial longitudinal magnetization acquired after the inversion pulse respectively.

### Generation of stable cells expressing rat *Oatp1a1* and human *Oatp1b3* transporters

Human embryonic kidney cells (HEK 293T), human triple negative breast cancer cells (MDA-MB-231), and rat glioma cells (C6) were obtained from a commercial supplier (American Type Culture Collection; ATCC, Manassas, Virginia, USA) and cultured in Dulbecco’s Modified Eagle Media (DMEM, Wisent Inc., Saint-Jean-Baptiste, Quebec, Canada) supplemented with 10% fetal bovine serum at 37°C and 5% CO_2_. Cells were routinely tested for mycoplasma using the MycoAlert mycoplasma detection kit (Lonza Group, Basel, Switzerland). Lentiviral production involved third-generation packaging and envelope-expression plasmids (pMDLg/pRRE, pRSV-Rev, and pMD2.G, Addgene plasmids: #12251, #12253, and #12259, respectively; gifts from Didier Trono). Lentiviral transfer plasmids were synthesized to encode *tdTomato* and rat-derived *organic anion-transporting polypeptide 1a1* for preclinical liver imaging or *zsGreen1* and human-derived *organic anion-transporting polypeptide 1b3* to assess clinical translation of MnTriCP-PhOEt, all under regulation of the human elongation factor 1 alpha promoter (pEF1α). To produce *Oatp1a1*- and *Oatp1b3*-encoding lentiviruses (LV-Oatp1a1 and LV-Oatp1b3, respectively), packaging, envelope and the relevant transfer plasmid were co-transfected into HEK 293T cells using Lipofectamine 3000 according to the manufacturer’s lentiviral production protocol (Thermo Fisher Scientific Inc., Waltham, Massachusetts, USA). Lentivirus-containing supernatants were harvested 24h and 48h post transfection, filtered through a 0.45-μm filter, and stored at −80°C prior to use. Rat C6 cells were transduced with lentivirus encoding *tdTomato* and rat-derived *organic anion-transporting polypeptide 1a1* (LV-tdT/FLuc/Oatp1a1) overnight in the presence of 4- to 8-μg/mL polybrene. Human MDA-MB-231 cells were transduced with lentivirus encoding *zsGreen1* and human-derived *organic anion-transporting polypeptide 1b3* (LV-zsG/Oatp1b3) following the same protocol. Transduced cells were washed, collected, and sorted using a FACSAria III fluorescence-activated cell sorter (BD Biosciences, Mississauga, Ontario, Canada) for either tdTomato and/or zsGreen1 fluorescence.

### Measuring relaxation rates of Oatp1-expressing cells incubated with MnTriCP-PhOEt

Non-transduced and Oatp1a1-expressing C6 cells, and non-transduced and Oatp1b3-expressing MDA-MB-231 cells were each seeded in T-175 flasks (2×10^6^) and grown for 3 days. Cells were trypsinized and suspended in 2 mL of DMEM containing either 1 mM Gd(III)-EOB-DTPA (Eovist/Primovist®, Bayer Health Care Pharma, Berlin, Germany), 1 mM Gd(III)-DTPA (Magnevist®, Bayer Schering Pharma, Berlin, Germany), or 1 mM MnTriCP-PhOEt for 60 minutes. Cells were centrifuged and washed three times with DMSO (30% v/v) in PBS to remove contrast agent remaining in the extracellular compartment, after which, 3×10^7^ cells were pelleted into 0.2-ml tubes, and then placed into a 1% agarose phantom. Inversion recovery images were acquired through a cross section of the tubes in the agarose phantom to measure spin-lattice relaxation rates of each cell condition, using the imaging parameters described above.

### *In vivo T*^1^-weighted imaging and *r*_1_ mapping in mice at 3 Tesla

All animal experiments were performed in compliance with an approved protocol of the University of Western Ontario’s Council on Animal Care (Animal Use Protocol 2016-026) and in accordance with the standards of the Canadian Council on Animal Care. Female nude mice (n=9 total, NU-*Foxn1*^nu^ strain; Charles River Laboratories, Wilmington, Massachusetts, USA) were anesthetized with 1%–2% isoflurane by using a nose cone attached to an activated carbon charcoal filter for passive scavenging, and positioned in a lab-built tray that was warmed to 40°C during MR imaging. For all mice, a pre-contrast image was acquired, followed by a slow rate (50 μL per minute, 500 μL volume total) intraperitoneal injection of 0.025 mmol/kg MnTriCP-PhOEt dissolved in 30% v/v DMSO (30% v/v) in saline and imaged for 60 minutes. All images were acquired on a clinical 3-Tesla GE MR750 clinical scanner (General Electric Healthcare, Milwaukee Wisconsin, USA), using a custom-built insert gradient [inner diameter =17.5 cm, gradient strength = 500 mT/m, peak slew rate = 3000 T/m/s, and a bespoke 3.5-cm diameter, 5.0-cm length birdcage radiofrequency coil (Morris Instruments, Ottawa, Ontario, Canada). *T*_1_-weighted images were acquired (n=4) using a three-dimensional Fast Spoiled Gradient Echo (FSPGR) pulse sequence: frequency FOV = 4.0 cm, phase FOV = 2.6 cm, slice thickness = 0.4 mm, TR = 12.0 ms, TE = 3.2 ms, flip angle = 60°, matrix size = 128×128, number of excitations = 1, receiver bandwidth = 31.25 kHz, acquisition time = 3:52 per image. To obtain *R*_1_ maps of kinetic data (n=2), the driven equilibrium single pulse observation of *T*_1_ (DESPOT1) approach^[37]^ was employed, using a 5° and 20° flip angle pair in an FSPGR pulse sequence: frequency FOV = 4.0 cm, phase FOV = 2.6 cm, slice thickness = 0.4 mm, TR = 6.9 ms, TE = 2.4 ms, matrix size = 100×100, number of excitations = 5, receiver bandwidth = 31.25 kHz, acquisition time = 3:39 per image. Pixelwise *in vivo* spin-lattice relaxation time maps for each timepoint were computed via MatLab (R2020a, MathWorks, Natick, Massachusetts, United States) by linking consecutive pairs of variable flip angle images and calculating the slope of their signal intensities according to the following equation:

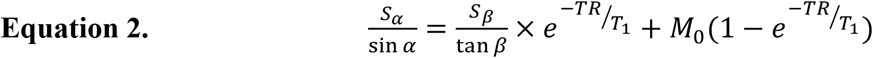

Where *S*_*α*_ is the FSPGR signal intensity associated with flip angle *α*, *S*_β_ is the FSPGR signal intensity associated with flip angle *β*, and *M*_0_ is the proportionality constant relating to the equilibrium longitudinal magnetization. Plotting 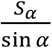 against 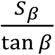 for each pixel allows for the determination of *T*_1_ from the slope *m* of this line as:

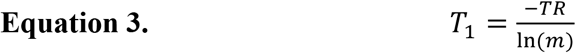

All resultant images were reconstructed and analyzed on open-source ITK-SNAP (Version 3.8.0) software^[38]^. Manual segmentation was performed for each mouse to create volumes-of-interest for the liver, both kidneys, and the heart, which were applied across pre-contrast and all post-contrast timepoints for each mouse to generate dynamic signal intensity (arbitrary units) or *T*_1_ time (ms) curves. Mean signal intensity ± standard deviation or mean *T*_1_ time ± standard deviation was then calculated for statistical testing.

### *Ex vivo* nuclear magnetic relaxation dispersion

To validate *in vivo R*_1_ calculations, a separate cohort of mice (n=3) were intraperitoneally injected with 0.025 mmol/kg MnTriCP-PhOEt dissolved in 30% v/v DMSO (30% v/v) in saline, or an equivalent volume of 30% v/v DMSO (30% v/v) in saline as a control. Forty-four minutes post-injection, mice were euthanized via isoflurane overdose and liver, kidney and muscle tissue were immediately harvested, gently rinsed with and placed in NMR tubes for low-field NMRD measurements. Spin-lattice relaxation rates (*R*_1_; Hz) ± error were acquired for each organ at the following field strengths: 0.0154023, 0.0280061, 0.0509018, 0.0925129, 0.1681517, 0.3056599, 0.555566, and 1.0008455 T using a fast field-cycling NMR relaxometer (SpinMaster FFC2000 1T C/DC, Stelar, s.r.l., Mede, Italy) as described above. *R*_1_ rates at each field strength were then averaged to determine statistical differences between MnTriCP-PhOEt and saline control mice.

### Statistics

Unless otherwise stated, all statistical analyses were carried out using Graphpad Prism software (Version 9.00 for Mac OS X, GraphPad Software Inc., La Jolla, California, USA, www.graphpad.com). For NMRD profiles, cubic spline regression curves were calculated with 4 knots and 4960 output segments. One-way Analysis of Variance (ANOVA) was performed followed by Tukey’s post-hoc multiple comparisons to determine statistical differences in longitudinal relaxation rates *in vitro*, *in cellulo*. and *in vivo*. For all tests, a nominal p-value less than 0.05 was considered statistically significant.

---

**Supplementary Figure 1.**
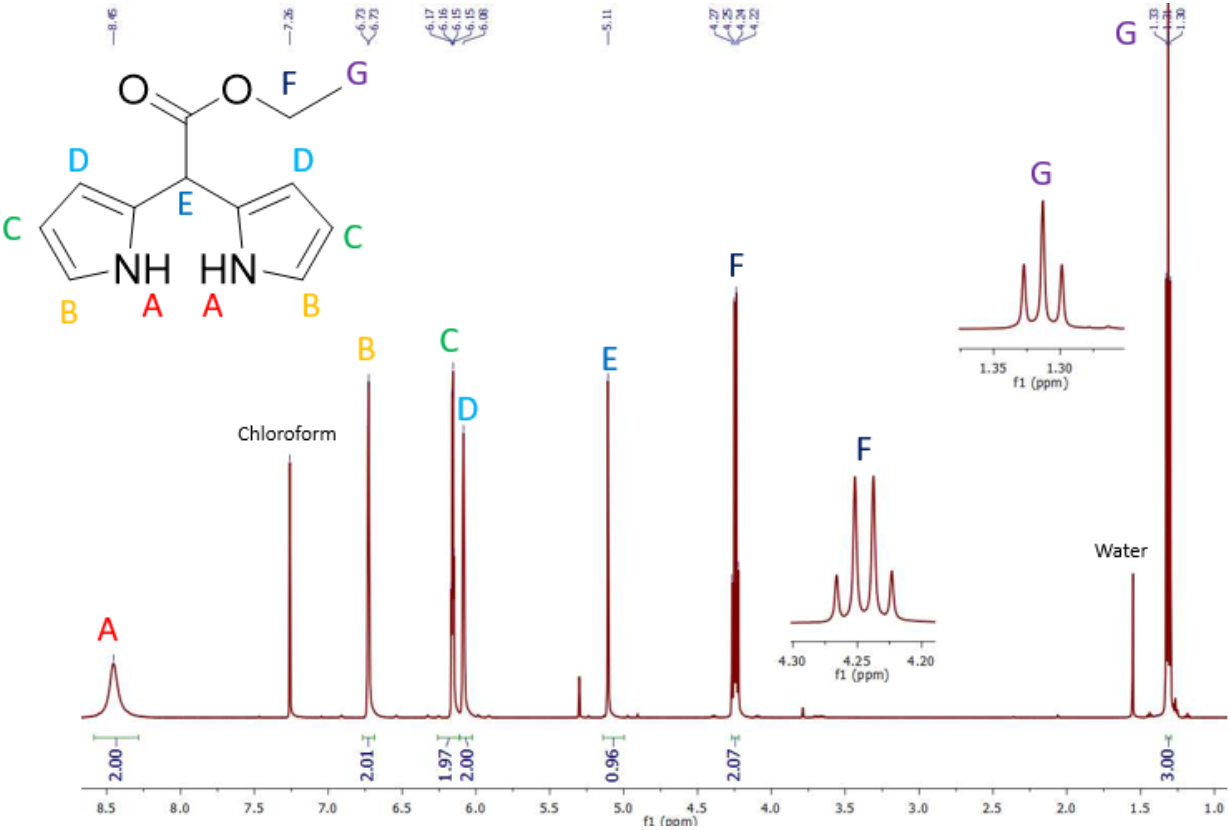
^1^H-NMR of EDP, a precursor to Mn(III)TriCP-PhOEt, in CDCl_3_.

**Supplementary Figure 2.**
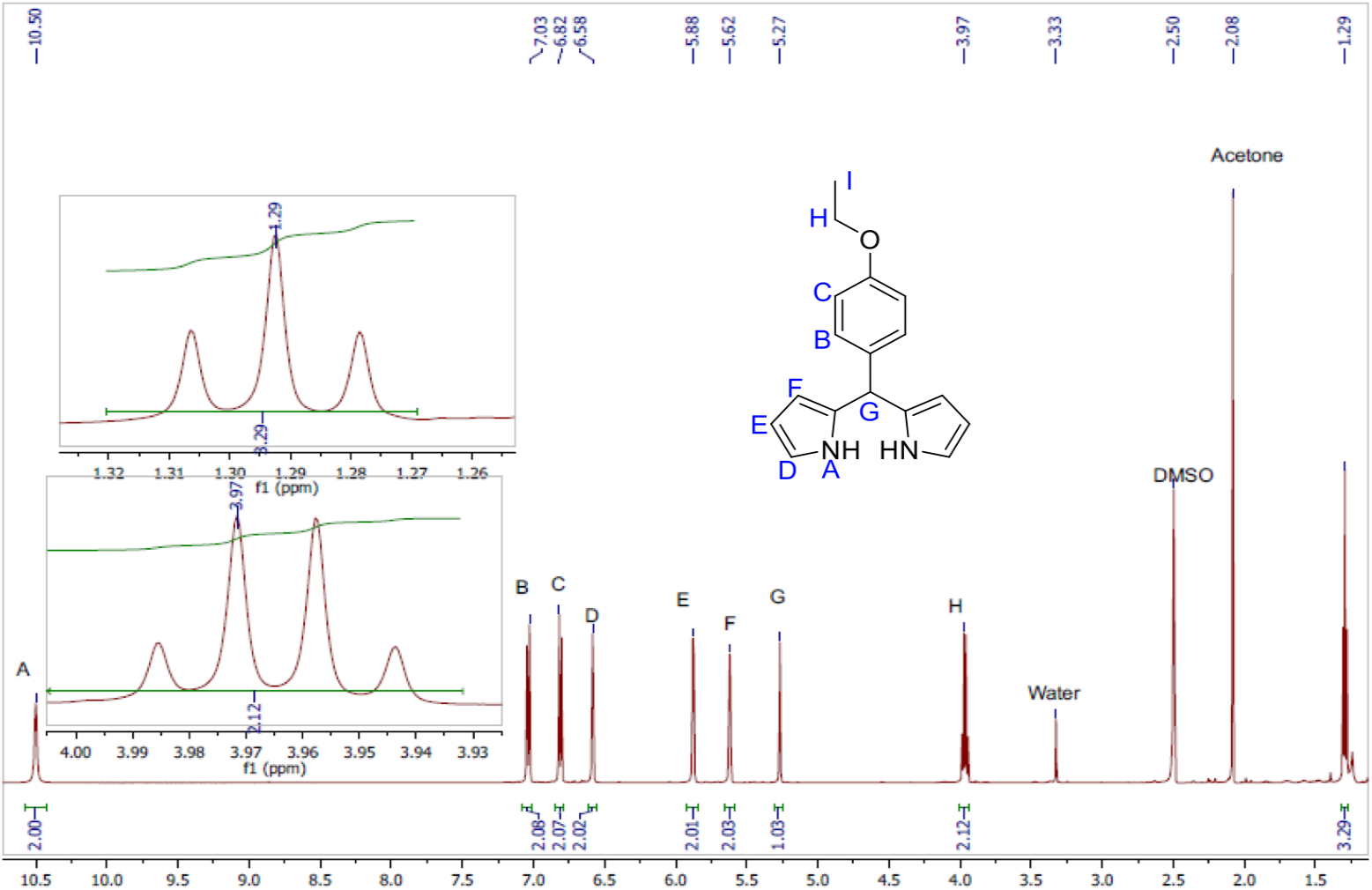
^1^H-NMR of DPM-PhOEt, a precursor to Mn(III)TriCP-PhOEt, in DMSO-d_6_.

**Supplementary Figure 3.**
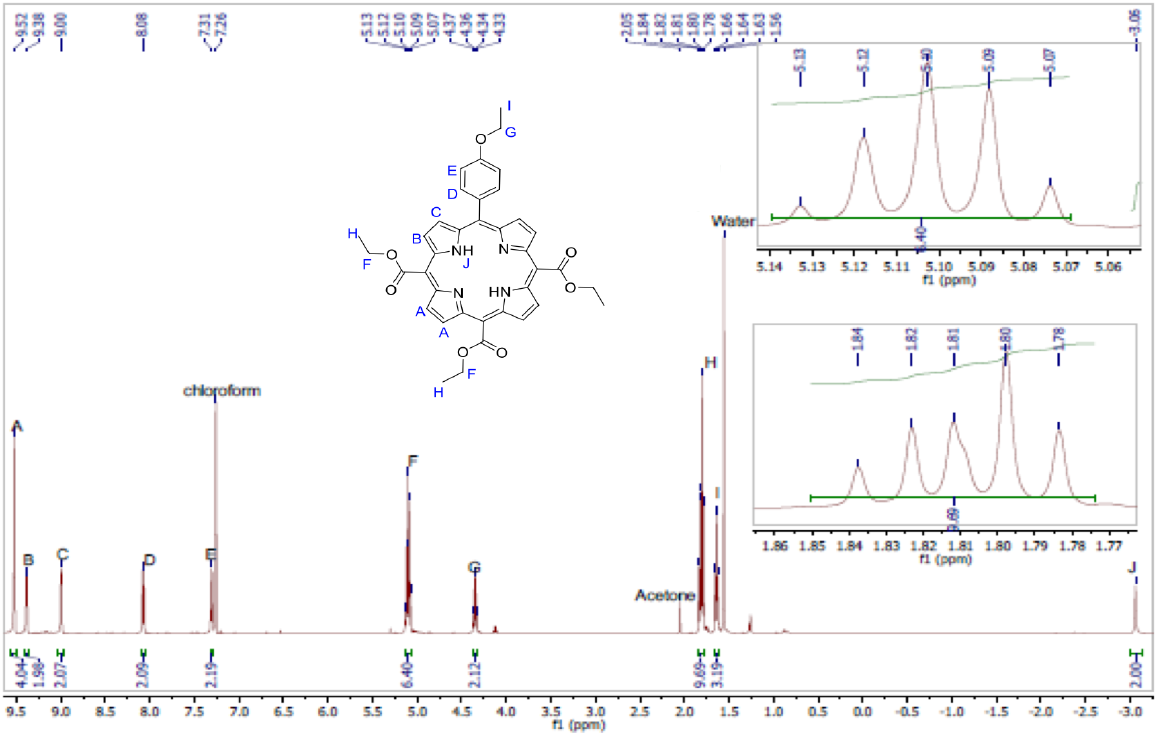
^1^H-NMR of TriEP-PhOEt, a precursor to Mn(III)TriCP-PhOEt, in CDCl_3_.

**Supplementary Figure 4.**
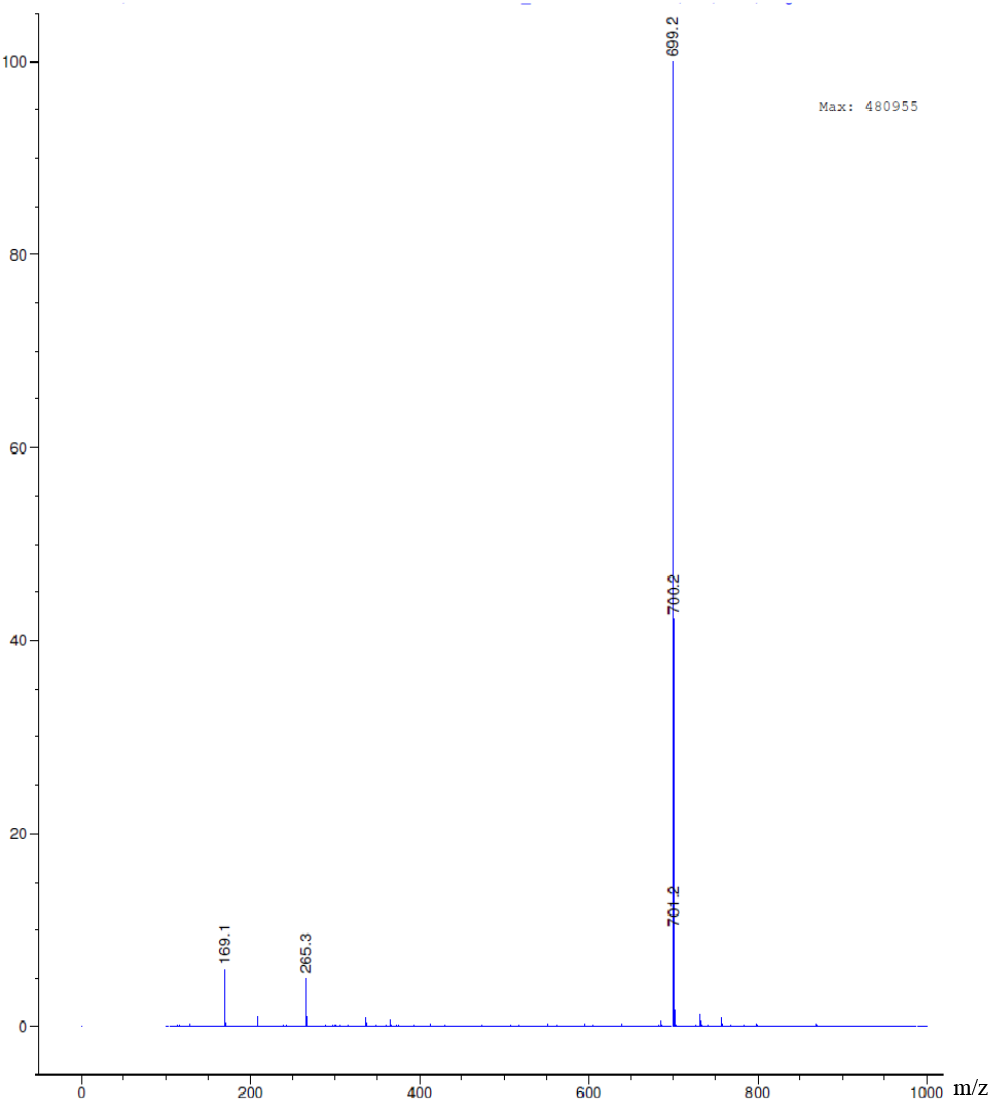
ESI-MS of Mn(III)TriEP-PhOEt, positive mode. ESI-MS found *m/z* = 699.2 ([M]^+^) Calculated for C_37_H_32_MnN_4_O_7_^+^, *m/z* = 699.17.

**Supplementary Figure 5.**
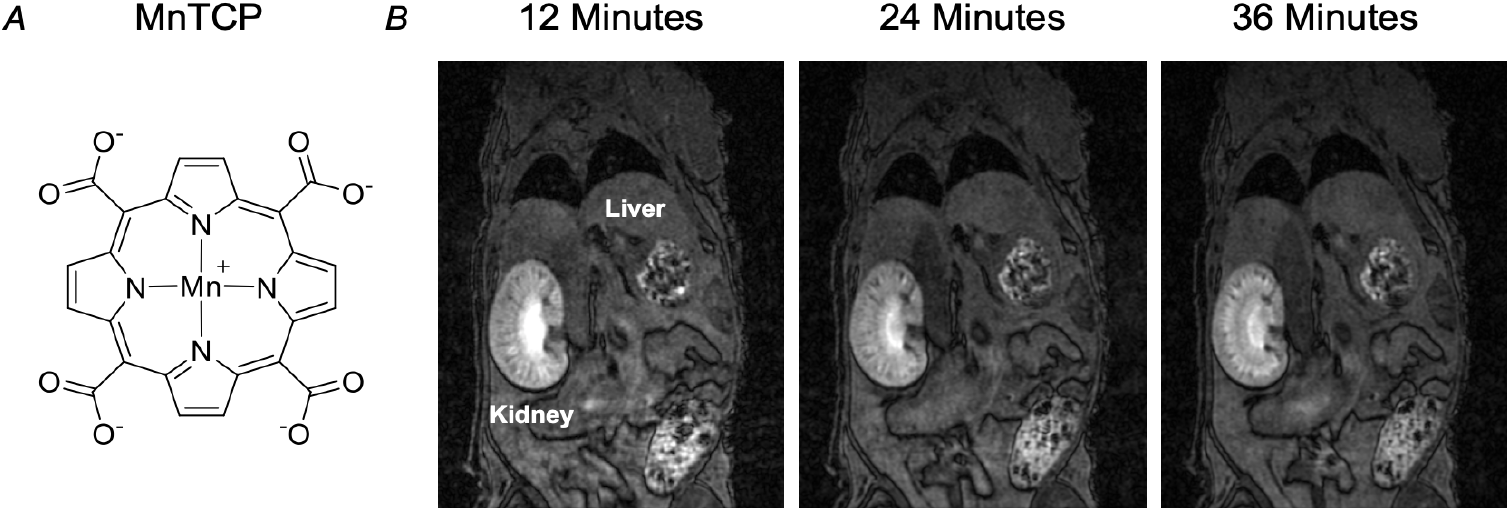
*In vivo* biodistribution of non-targeted manganese(III) porphyrin. A, Chemical structure of manganese(III) porphyrin-5,10,15,20-tetracarboxylate (MnTCP). B, *T*_1_-weighted images of a female mouse acquired 12, 24, and 36 minutes post intravenous administration of 0.1 mmol/kg Mn(III)TCP at 3 Tesla.

